# DIY-NAMIC behavior: A high-throughput method to measure complex phenotypes in the homecage

**DOI:** 10.1101/2020.04.24.059980

**Authors:** Jun Ho Lee, Selin Capan, Clay Lacefield, Yvonne M. Shea, Katherine M. Nautiyal

**Author notes:** To whom correspondence should be addressed: Katherine M. Nautiyal, PhD, 6207 Moore Hall, Hanover, NH 03755.

## Abstract

Complex behavioral assessment is becoming increasingly necessary in order to comprehensively assess *in vivo* manipulations in rodent models. Using operant behavioral paradigms provides rich data sets allowing for the careful analysis of behavioral phenotypes. However, one major limitation in these studies is the expense and work-load that are required using traditional methods. The equipment for commercial operant boxes can be prohibitively expensive, and the daily experimenter effort and mouse costs required for these studies is extensive. Rodents are generally trained on task-specific paradigms for months, tested every day for 5-7 days per week. Additionally, appetitive paradigms usually require food restriction and are also commonly run in the non-active light phase of the rodent circadian rhythm. These limitations make operant behavioral testing especially difficult during adolescence, a time period of interest with regards to the development of adult-like phenotypes and a high-risk period for the development of neuropsychiatric disorders, including those which involve impulsive behavior. In order to address these issues, we developed an automated, inexpensive, open-source method which allows the implementation of most standard operant paradigms in the homecage of rodents in shorter time frames without food restriction, and with much less experimenter effort. All construction and code for the DIY Nautiyal Automated Modular Instrumental Conditioning (DIY-NAMIC) system are open source. We demonstrate their utility here by measuring impulsive behavior in a pharmacology experiment, as well as in adolescent mice.

**Significance statement:** Rigorous behavioral assessment is critical to understand the neural basis of neuropsychiatric disorders using animal models. Operant behavioral paradigms provide the ability to measure complex phenotypes, however, traditional methods generally require time-consuming daily training for many weeks. We designed, built, and tested an open-source automated homecage system for appetitive instrumental conditioning that enables testing in shorter timeframes with reduced experimenter effort.

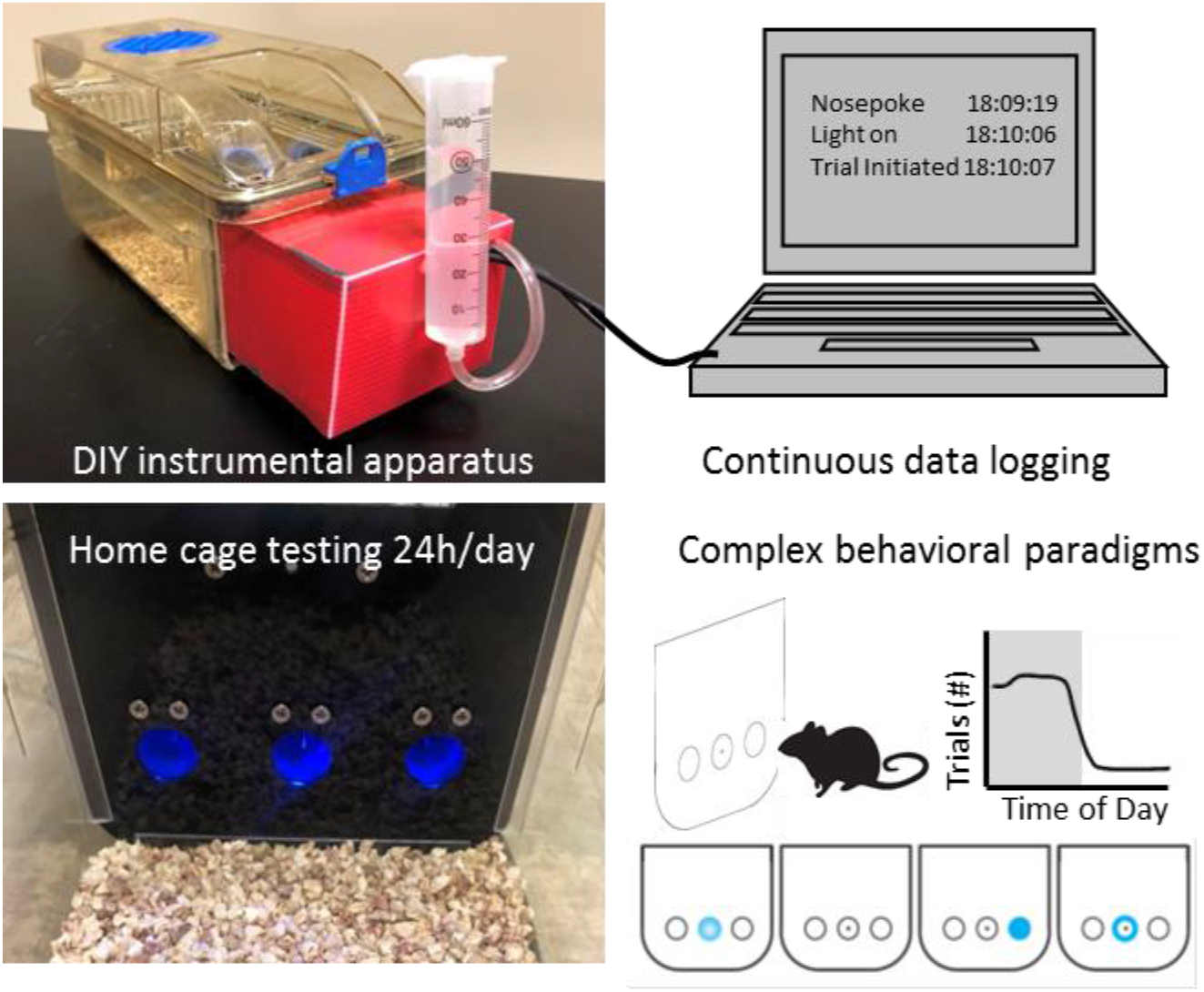

## Introduction

Complex behavioral phenotyping is critical for understanding the effects of neural and genetic manipulations in animal models. In fact, even funding agencies are requesting the use of more multi-dimensional behavioral measures in neuroscience research, especially in animal models for neuropsychiatric disorders (Bale et al., 2019; NIMH, 2019). Operant behavioral paradigms provide an excellent framework to assess multiplexed behavioral effects of biological manipulations in rodent models. However, traditional methods generally take months to run, require daily training, prolonged food restriction, and involve costly equipment. An ideal testing system would eliminate daily experimenter intervention and minimize experiment time-lines by allowing hundreds of trials per day, while also removing the need for food deprivation, and allowing testing and data logging automatically 24h/day. An automated homecage operant system would also provide more reproducibility given reduced experimenter and protocol-introduced variability.

Impulsive behavior is found in many neuropsychiatric disorders such as substance use disorder, behavioral addictions, and attention deficit hyperactivity disorder (Jentsch, 2008; Winstanley, 2011; Bari and Robbins, 2013). Importantly, there is a long history of careful behavioral dissection of impulsivity using operant paradigms in rodent models (Dalley et al., 2011; Bari and Robbins, 2013). There have been many studies focused on characterizing various aspects of impulsive behavior, and assessing related phenotypes. One distinction that has been made is between different types of impulsivity including impulsive choice (e.g., delaying gratification), versus impulsive action (e.g., withholding responding) (MacKillop et al., 2016; Nautiyal et al., 2017). While both are frequently elevated in psychiatric disorders presenting with disordered impulsivity, they can be statistically and biologically dissociated as separate (and uncorrelated) components in the pathogenesis of many disorders (Winstanley et al., 2006; Brevers et al., 2012; Grant and Chamberlain, 2014). Our focus on measuring impulsive action in mouse models to understand its neural basis has required extended daily task-specific training in operant paradigms, taking 2 months or more, for example in the standard procedure for the 5 choice serial reaction time task (Humby et al., 2005; Bari et al., 2008; Fletcher et al., 2013). Furthermore, studying the development of impulsivity during adolescence has not been possible in standard operant paradigms for impulsive action, given that adolescence lasts less than 1 month in mice (spanning approximately 30-60 days postnatal) (Workman et al., 2013; Bell, 2018).

Studying the behavioral and neural development of impulse control during adolescence in mouse models is important for understanding the adolescent sensitive period as it relates to the etiology of a number of neuropsychiatric disorders (Paus et al., 2008; Casey, 2015; Schulz and Sisk, 2016). Specifically, animals show high impulsivity, novelty seeking, and risky behavior during adolescence, a potentially evolutionarily adaptive behavioral change which is important to promote independence at sexual maturity. Additionally, understanding the neural changes that lead to the decrease in these behaviors at the end of the adolescent period is also important for delineating the neural circuits that may be disordered when these phenotypes persist into adulthood, and may emerge as important factors in psychiatric disorders.

A large literature implicates serotonin in the regulation of impulsive behavior in humans and animal models (Brunner and Hen, 1997; New et al., 2001; Bevilacqua et al., 2010; Miyazaki et al., 2012; Stoltenberg et al., 2012; Nautiyal et al., 2015b). Pharmacological and genetic manipulations in rodents have been a large source of our knowledge on the neural mechanisms that drive impulsivity. Most of this preclinical testing has occurred in MedAssociates operant chambers using complex and well-validated operant paradigms (Robinson et al., 2008; Zeeb et al., 2009; Bari and Robbins, 2013). These studies have delineated the complex role of serotonin signaling in the differential modulation of a number of facets of impulsive behavior (Harrison et al., 1997; Winstanley et al., 2004a; Winstanley et al., 2004b; Dalley and Robbins, 2017). Global depletion of serotonin seems to increase impulsive action, with limited effects on impulsive choice (Harrison et al., 1997; Winstanley et al., 2004a). Additionally a number of the 14 different serotonin receptors have been implicated in the modulation of impulsive action including 5-HT_2A_ and 5-HT_2C_ receptors using pharmacology (Winstanley et al., 2004b; Fletcher et al., 2011), and 5-HT_1B_ and 5-HT_2C_ using genetic loss-of-function models (Brunner and Hen, 1997; Fletcher et al., 2013; Nautiyal et al., 2015a).

In order to measure impulsivity and a range of related behavioral parameters, we sought a more time- and cost–effective method for fine-grain measurement of behavior in rodents. In recent years, there have been a number of automated and open-source developments for complex behavioral testing which can increase throughput as well as data reproducibility (Nguyen et al., 2016; Ali and Kravitz, 2018; Godynyuk et al., 2019; Venniro et al., 2019; Luciani et al., 2020). ‘do-it-yourself’ (DIY) operant behavioral testing apparatus have been developed for use with standard session-based testing methods which provide relatively inexpensive and customizable apparatus making behavioral testing more accessible (Devarakonda et al., 2016). Additionally, there have been modifications made to traditional commercial equipment, and also to standard lengthy paradigms making shorter training times possible (Remmelink et al., 2017; Sasamori et al., 2018). What was still lacking, however, was a method that combined these advantages to allow for both homecage and continuous assessment of operant behavior, with liquid reward in an affordable open-source system.

Our goal was to design, build, validate, and disseminate a novel method which allows for continuous quantitative and robust operant behavioral testing. We developed the DIY Nautiyal Arduino Modular Instrumental Conditioning (DIY-NAMIC) system as an inexpensive, open-source method which allows mice to perform operant trials in their homecage to receive their daily consumption of water, in self-initiated trials. This promotes rapid acquisition of task-specific learning in behavioral paradigms given the high number of daily trials that mice perform each day. With continuous testing and automatic data-logging the DIY-NAMIC system enables high throughput data collection 24h/day, 7d/week. Using the DIY-NAMIC system, we were able to measure impulsive action as well as a number of other behavioral parameters in under three weeks, with only twice weekly (less than 1h long) experimenter effort. We show that mice rapidly acquire Pavlovian and stimulus-response associations, and basic cue-based discrimination learning within a week. Furthermore, we show that effects of pharmacological manipulations can be measured with the DIY-NAMIC system, and demonstrate that activating the serotonin 1B receptor (5-HT_1B_ R) reduces impulsive action. Finally, our studies illustrate the ability to test impulsivity in adolescent mice, a period of development which is difficult to test in mice using standard paradigms. Overall, the studies presented here validate the use of the DIY-NAMIC system for assessing multi-dimensional behavior including impulsivity. Furthermore, we present novel data showing serotonergic modulation of the neural basis of impulsive behavior, and by characterizing adolescent impulsivity in mice.

## Methods

### DIY-NAMIC System Overview

The DIY-NAMIC system was built for compatibility with most facilities’ standard rodent homecages in high-density ventilated racks by replacing one cage wall with a modular system that delivers stimuli and rewards, and detects and records behavioral responses (Fig 1A). It allows for the implementation of standard Pavlovian and operant-based tasks though with less effort, time, and cost. The DIY-NAMIC system is controlled by an Arduino microprocessor board and easily sourced electronics allowing for an inexpensive and customizable apparatus. The initial design used in these studies includes three solenoid-delivered liquid dispensing noseports, each with an infrared head-entry detector and LED cue light (Fig 1B). Mice do not undergo food restriction and the DIY-NAMIC system permits behavioral testing during the active phase of their circadian rhythm and eliminates daily, potentially stressful, handling. The system is designed for liquid delivery enabling fine control of volume (versus pellet delivery). The receptacles are shallow, allowing reward receipt with large head stages, such as those for miniature head-mounted microscopic imaging. Analysis scripts, written in Python, are available for use and modification as needed. Given their modular construction and open source development, the hardware for the DIY-NAMIC system is customizable and inexpensive to build.

**Figure 1.**
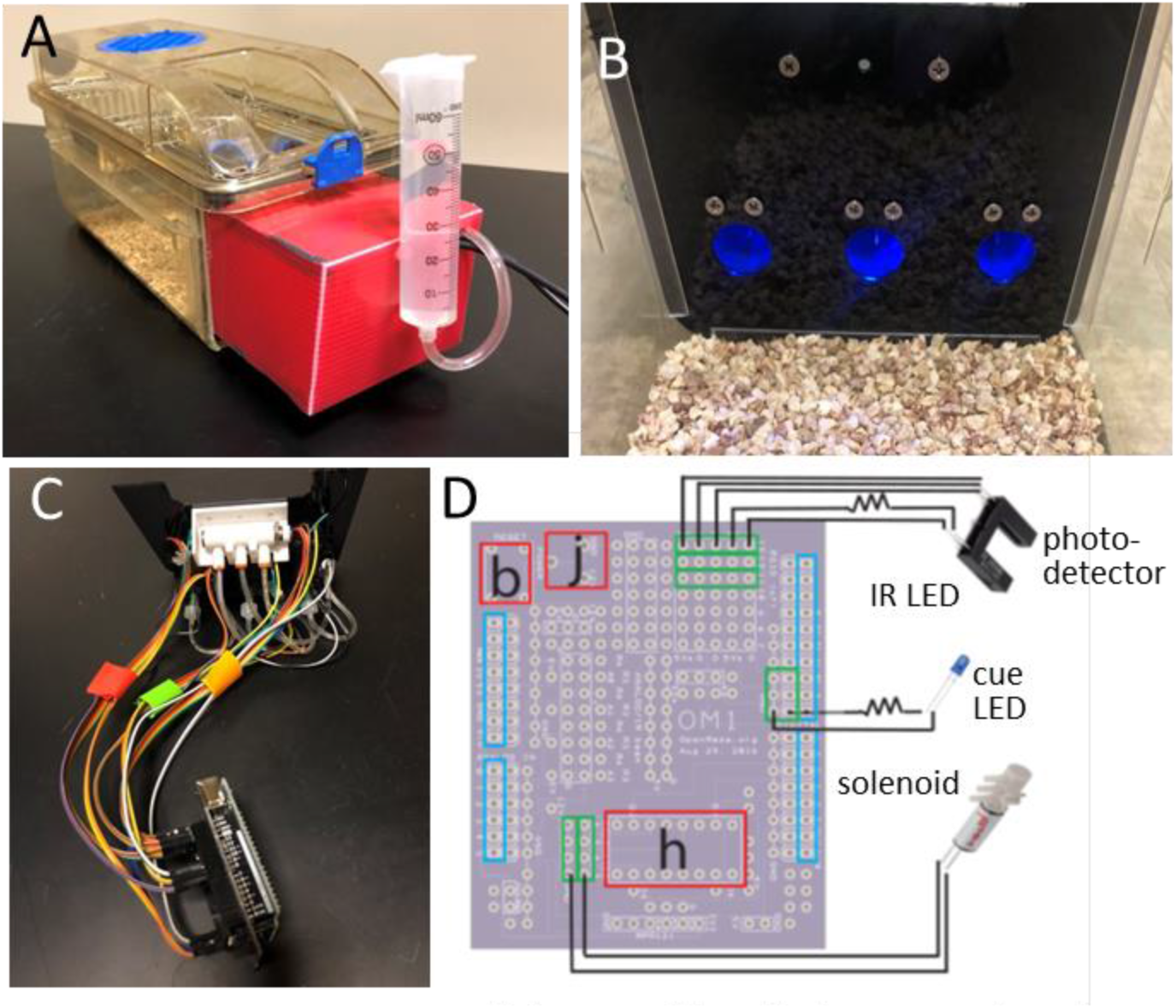
The DIY-NAMIC system is integrated into the homecage of standard mouse cages. (A) The Arduino and all components of the DIY-NAMIC system are enclosed within the red encasing with the exception of a syringe for water or liquid reward, mounted on the outside. Modified cages remain compatible with most standard high-density ventilated racks. (B) Inside homecage view of the DIY-NAMIC system including three nose ports with blue LEDs illuminated, each of which contain a spout for liquid reward. (C) The inside of the apparatus is shown including LEDs, solenoids, and IR head entry detectors wired to an Arduino UNO through an OpenMaze (OM1) shield. (D) Schematic wiring diagram shows the wiring of one set of components (for one noseport of three total) to the OM1 shield. Blue rectangles indicate location of stacking headers for connection to Arduino. Red rectangles indicate locations of chip socket for H-bridge (h), push button for flush (b), and barrel jack (j). Green rectangles indicate location of female headers for connection to IR (LED and photodetector), cue LED, and solenoid components.

### DIY-NAMIC Build Instructions

All materials required to build DIY-NAMIC boxes are easily sourced from common websites with the total cost of all parts required for each box at less than $300 (Table 1). All build instructions, detailed diagrams, and design files are located online at www.GitHub.com/DIY-NAMICsystem. We estimate that a user familiar with soldering and the basics of electronics and Arduino functions could build 8 cages within a week, in an assembly-line approach (Fig 1C). Standard mouse ventilated home cages (5.5” × 14” × 5”, Techniplast) were modified by cutting one short wall off with a rotary tool, and U channels were affixed with Loctite 401 inside the cut edge. A plexi-glass wall was laser cut (ordered from Ponoko; design file available online) to fit the dimensions of the cage, and slotted into the U channels. The wire and ventilated cage tops fit as usual. Three nose ports per cage were 3D printed (Shapeways, design file available online) and attached to the plexi-glass wall with flat head screws. The leads of the LEDs (5mm ultra bright diffused round blue light) used for cue lights were threaded through the top holes in the back of the noseport, and a current-limiting resistor (1 kOhm) was soldered to the anode. A female Dupont connector on a female to male 12in jumper wire was attached to each of the cathode and resistor ends and soldered in place. Photo interrupters (IRs) were cut in half to separate the infrared LED and phototransistor and each part was inserted into casing on either side of the designed noseport, and a hex nut was threaded onto the screws affixing the noseports to the plexiglass which secured the IRs in place. A resistor was wired to the LED of the IR, and female Dupont connectors of jumper wires were attached to each of 5 IR leads, as described for the cue LEDs. Large gauge hypodermic needles (21Ga) were blunted by removing the beveled sharp tip with a rotary tool and then sanded smooth with a belt sander. The needles were inserted into the lower hole on the back of each nose port, and secured with hot glue. Luer locks connected the needle to tubing (1/16” ID), which connected to the middle port of the solenoid valve (3-ported, 5V-30PSI, Lee Company). The two solenoid leads were attached to female Dupont connectors of jumper wires. The three solenoids (one per noseport) were connected to a 3-way manifold from the solenoid port distal to the leads, with the same tubing (1/16” ID). A small closed piece of tubing was attached to the proximal solenoid port as a cap. The manifold was fed water from a 60ml syringe via tubing (1/4” ID)

**Table 1.**

| Component | Specification | Supplier/<br>Part ID | Cost | URL |
| --- | --- | --- | --- | --- |
| Noseports | 3D printed with black natural versatile plastic (3 needed/cage) | Shapeways | \$14.10 | File:<br>Printing:<br>https://www.shapeways.com/ |
| Plexiglass wall* | Black Acrylic, 1.5mm thick | Custom laser cut at | \$6.30 | File:<br>Laser cutting: |
|  |  | Ponoko |  | https://www.ponoko.com/laser-cutting |
| U channel | Clear impact-resistant polycarbonate U-channels (0.235" wide, 0.3" high) | McMaster-Carr 1753K61 | \$4.06/4ft | https://www.mcmaster.com/catalog/126/3817 |
| Screws | Philips Flat Head: 4-40 Thread Size, 5/16" Long (6 needed/cage) | McMaster-Carr 91771A107 | \$4.53/100 | https://www.mcmaster.com/catalog/126/3227 |
| Photo Interrupter | GP1A57HRJ00F (3 needed/cage) | Sparkfun 09299 | \$2.50 | https://www.sparkfun.com/products/9299 |
| LEDs | 5mm LED Ultra Bright Diode, DC 3V 20mA 0.06W Diffused Round Blue light (3 needed/box) | Amazon | \$6.00/100 | https://www.amazon.com/gp/product/B06XFVBQDY/ref=ppx_yo_dt_b_search_asin_title?ie=UTF8&psc=1 |
| Resistors | 1 K Ohm (6 needed/cage) | Sparkfun | \$0.95/20 | https://www.sparkfun.com/products/14492 |
| Needles | BD Precision Glide Needles (21G, 0.8 mm x 50 mm; 3 needed/cage) | Fisher Scientific 14-821-13N | \$29.22/100 | https://www.fishersci.com/shop/products/bd-precisionglide-single-use-needles-regular-bevel-regular-wall-10/1482113n |
| Luer Lock | Masterflex fitting, Male luer lock to hose barb adapter (3 needed per cage) | Cole-Parmer UX-45504-19 | 16.20/pkg 20 | https://www.coleparmer.com/i/masterflex-fitting-polycarbonate-straight-male-luer-lock-to-hose-barb-adapter-1-4-id-25-pk/4550419?searchterm=UX-45504-19 |
| Tubing 1 | Tygon E3603 Laboratory Tubing ID 1/16", OD 1/8", Wall 1/32" | Component Supply TET-062A | \$0.32/ft | https://componentsupplycompany.com/product-pages/tygon-e-3603-lab-tubing.php |
| Solenoid | LHDA0533115H HDI PORTED 3P-5V-30PSI (3 needed/cage) | Lee Company | \$63.10 | https://www.theleeco.com/products/electro-fluidic-systems/solenoid-valves/control-valves/lhd-series/3-port/ported/ |
| Tubing 2 | Tygon E3603 Laboratory Tubing | Component Supply | \$0.77/ft | https://componentsupplycompany.com/product-pages/tygon- |
|  | ID 1/4", OD 5/16", wall 1/32" | TET-250A |  | e-3603-lab-tubing.php |
| 3-way manifold | Push-to-connect Tube Fitting for Air and Water, 3 outlet manifold, 1/8 NPT Male x 1/8" Tube OD | McMaster Carr 5203K929 | \$12.66 | https://www.mcmaster.com/catalog/126/239 |
| Syringe | 60ml syringe, Luer-lock | Fisher Scientific 14-955-461 | \$32.54 /50 | https://www.fishersci.com/shop/products/sterile-syringes-single-use-12/14955461?keyword=true |
| Arduino | Arduino Uno Rev3 | Arduino 80583334 90090 | \$22.00 | https://store.arduino.cc/usa/arduino-uno-rev3 |
| Shield | OM1 shield OpenMaze.org | Oshpark OM1 board v082916 | \$28.05 /3 | https://oshpark.com/shared_projects/ea7NpG7p |
| IC socket | 16-pin 0.3" Chip | Adafruit 2203 | \$0.95 | https://www.adafruit.com/product/2203 |
| H-Bridge | Dual H-Bridge Motor Driver – 600mA L293D | Adafruit 807 | \$2.95 | https://www.adafruit.com/product/807 |
| DC barrel jack | Breadboard-friendly 2.1mm DC barrel jack | Adafruit 373 | \$0.95 | https://www.adafruit.com/product/373 |
| Jumper wires | 12" female to male jumper wires | Sparkfun 09386 | \$34.95 /100 | https://www.sparkfun.com/products/9386 |
| Headers | Stacking headers with standard 2.54mm spacing | Amazon | \$8.09 / 40 | https://www.amazon.com/gp/product/B07JN1KP7W/ref=ppx_yo_dt_b_search_asin_title?ie=UTF8&psc=1 |
| Heat Shrink Kit | 1/8" diameter | Sparkfun 09353 | \$7.95/ 95 pieces | https://www.sparkfun.com/products/9353 |
| Power supply | 9V DC 1000mA, AC 100V-240V | Adafruit 63 | \$6.95 | https://www.adafruit.com/product/63 |

\*This design is based on dimensions of the cages used in our facility's high density ventilated rack system, and would need to be modified to fit cages used in other facilities.
Other general tools and supplies used: Rotary Tool (Dremel), Loctite 401, soldering iron (Weller WES51 Analog Soldering Station, Sparks, MA); Surge Protector; Multi-port USB hub

An Arduino UNO R3 board was connected via double stacking header pins to an Arduino “shield” from OpenMaze.org (OM1 board). Wiring is documented in detail on the circuit diagram and the build instructions online.. Specifically, 32 female stacking headers were inserted and soldered into the printed circuit board (OM1 board; see Fig 1D) to connect it to the 32 Arduino Pinouts. An integrated circuit socket for an H-bridge and DC Barrel Jack were also soldered to the OM1 board to provide additional power required for the solenoids. A push button, used for a system water flush, was also soldered to the OM1 board. Female headers were soldered to the OM1board to allow for the simple ‘snap-in’ connection of all of the components via the male ends of the jumper cables wired to the LEDs, solenoids, and IR detectors, as described above (Fig 1D). A wall power supply was connected to the board through the DC barrel jack connector. Arduinos were connected via 10-USB hubs to a PC to allow for continuous data logging of 16 cages directly to a PC using Processing software. A test program is provided online which allows for the verification of correct functioning and troubleshooting of all of the components. Additionally, when the data collection is started through Processing, the data file reports correct initiation of the IRs (not blocked) to validate their correct functioning.

### Mice

Mice were bred in the vivarium at Dartmouth College. Mice were weaned at postnatal day (PN) 21 into cages of 2-4 same sex littermates and maintained on *ad libitum* chow and water in a 12:12 light-dark cycle. Corncob bedding was used instead of shavings in the homecage to reduce the likelihood of occlusion of noseports or liquid ports. For the initial characterization of behavior shown, male and female mice (N=9) were tested in DIY-NAMIC boxes beginning at 14 weeks of age. For the study of impulsivity in adolescents, male (N=9) and female (N=9) mice were placed individually in DIY-NAMIC boxes at 45-47 days postnatal (Adolescents, N=9) or 14-15 weeks postnatal (Adults, N=9). The pharmacology study was performed on mice aged 11-20 weeks old (N=18). All mice were placed in DIY-NAMIC boxes individually, with environmental enrichment including a plastic igloo and nesting material with *ad libitum* chow, with the water bottle removed. All mice were on a mixed C57/Bl6J and 129Sv/Ev background. All procedures were approved by the Dartmouth College Institutional Animal Care and Use Committee.

### Behavioral Training

Mice were singly housed and initially habituated to the DIY-NAMIC boxes and to retrieving 10μl water reward from the center noseport delivered by a solenoid. Based on the specifications of our build, including the height of the syringe, we empirically determined the volume to time relationship for size of the water reward dispensed relative to how long the solenoid was open (volume = 0.156*time-0.00134; r^2^=0.999). Opening the solenoid for 72ms resulted in 10μl of water dispensed, with a 3.4% coefficient of variation. During the first three days, a reward retrieval training paradigm (“P1”; Fig 2A) was presented in which the cue light in the center port was illuminated and mice received 10μl of water upon poking into the center port. After water retrieval, a variable inter-trial interval (ITI), with an average of 45s preceded the next light illumination.. The ITI remained at 45s average, and the water reward volume remained at 10μl, for all paradigms

**Figure 2.**
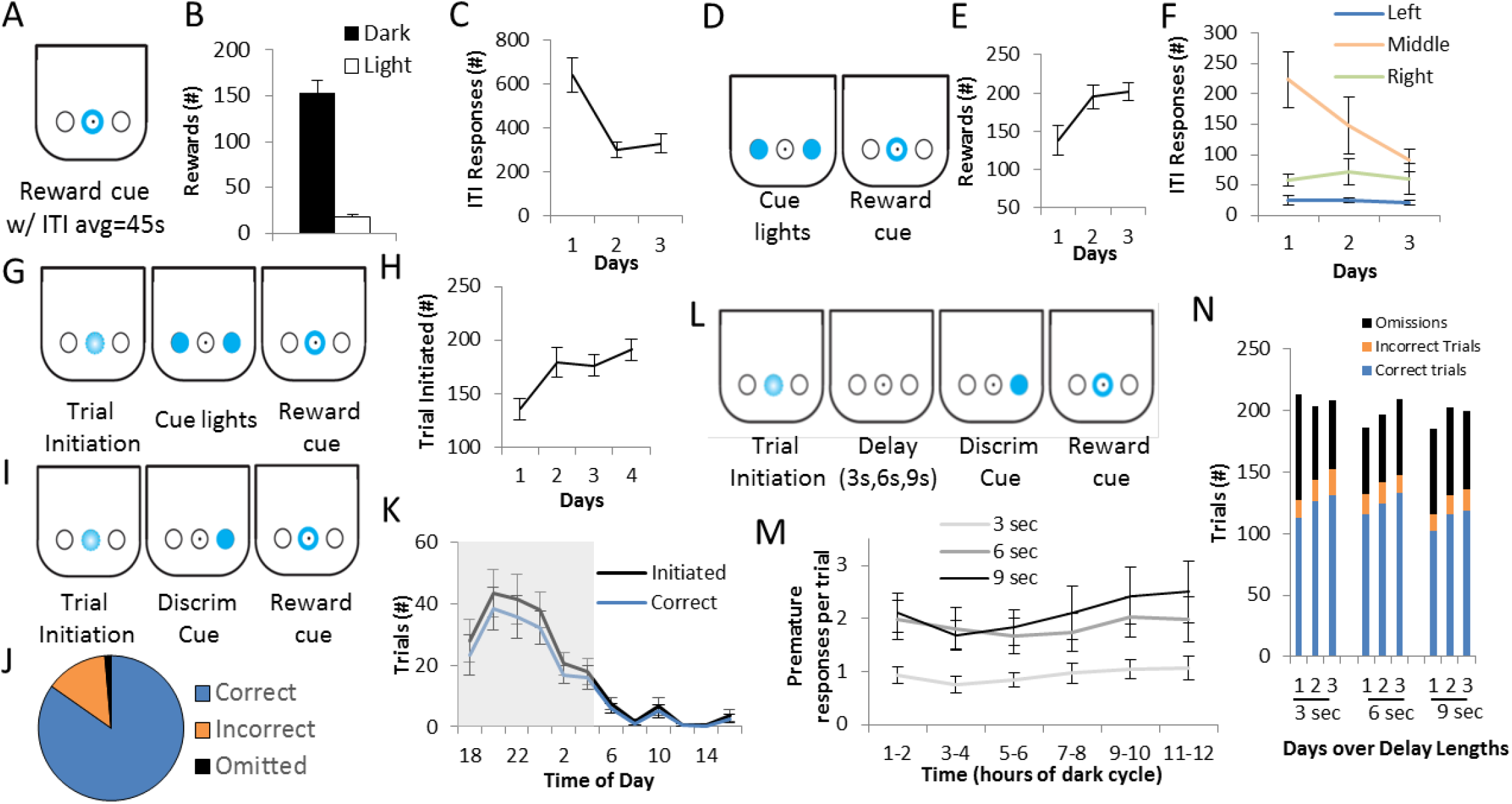
Measuring operant behavior with the DIY-NAMICsystem. A) Reward retrieval training (PI) occurred with presentations of reward available in trials with an ITI average of 45sec. Schematic shows the center reward port is illuminated when the reward is available. B) The number of rewards retrieved over 3 days is shown, separated by light and dark cycle. C) Number of pokes during the ITI when the reward is unavailable is shown over 3 days. D) Schematic for continuous reinforcement training (P2) shows that both side ports are illuminated during a trial, and pokes to either results in reward availability. E) The number of rewards received is shown across the 3 days of training. F) ITI responding is shown over 3 days for each noseport. G) Schematic for P3 which includes trial initiation shows how trials become self-initiated by a poke to the blinking center port. H) Number of self-initiated trials over X days of the paradigm. I) Schematic describes the operant cue discrimination paradigm (P4), in which the correct/rewarded port was illuminated by the LED. J) The group average shows that mice perform correctly on 85% of trials, and have relatively few incorrect (14%) and omitted (1%) trials. K) The number of trials initiated, and the number of correct trials, are shown across 24h, binned by 2h. The shaded gray area Indicates the dark phase of the light cycle. L) Schematic shows the modified 2-choice serial reaction time task which was used to assay impulsive action. M) Three-day averages of premature responses per trial over three delay lengths, shown by 2h bins over the dark cycle. N) Total number of correct, incorrect, and omitted trials on each of three crays run on three delay lengths. All group averages are shown for N=9 mice, with error bars representing standard error of the mean.

Subsequently, mice were trained on a continuous reinforcement schedule (“P2”; Fig 2D), for 3 days to nose poke in one of the two side nose ports when illuminated with a cue light. Pokes were rewarded in the center port. Both ports were illuminated with cue lights indefinitely on each trial, and a poke to either port was rewarded, and then the ITI followed. A trial initiation requirement was then added (“P3”; Fig 2G), and each trial began following a nosepoke to the center port blinking (1 Hz). Next, a basic cue discrimination requirement was added (“P4”; Fig 2I) in which only one of the two sides were illuminated, and only pokes to the illuminated port were reward. For 3 days, only correct nosepokes produced an outcome (reward) and incorrect pokes (pokes to the non-illuminated port) had no consequence. Subsequently, incorrect pokes resulted in trial termination and ITI onset, and trials were capped at 5s (no response before 5s resulted in ITI onset and the trial was considered an omission). After 2 days, the trial response duration was shorted to 1.5s. Finally, a modified 2-choice serial reaction time test paradigm (Fig 2K) was tested as a measure of impulsive action, by introducing a delay period between trial initiation and the cue light onset. The time delay was increased over 3 lengths: 3s, 6s, and 9s – with each presented for 3 days. Nose pokes during the delay were recorded but had no influence on the outcome of the trial (i.e. there was no timeout/punishment period or cue). Only a poke during the illuminated cue was necessary for a rewarded trial. All programs were switched during the light cycle, roughly 4-6h after light onset. Arduino programs for all paradigms are provided online including schematics illustrating trial structure and detailed input/output descriptions www.GitHub.com/DIY-NAMICsystem. The welfare of the mice was monitored by weight at least twice weekly. Data was processed regularly to ensure all mice were receiving adequate hydration from the system. Any mouse that did not receive more than 100 rewards/day for more than 2 days in a row, or lost more than 10% of their baseline bodyweight, was removed from the DIY-NAMIC cage and returned to *ad libitum* water. Two out of 47 mice had to be removed from the experiments.

### DIY-NAMIC Data Collection and Processing

Behavioral responses, as well as stimuli and reward presentations are logged with event and timecode stamps which are written directly into text files through Processing scripts. Following data collection, initial analysis is performed off-line using Python scripts in order to concatenate data log files, and extract relevant metrics including total number of rewards, and total number of nosepokes to each port, and nosepokes to each port during the ITI, reward presentation, cue presentation, and delay period. Premature pokes were considered pokes to one of the two side ports during the delay period. Time bounds are input to allow data extraction in desired time bins (e.g., minutes, hours, dark cycle, or days). All data collection and processing scripts are available online at www.GitHub.com/DIY-NAMICsystem.

### DIY-NAMIC Maintenance

Water is flushed through the system at a minimum of every 2 weeks, by pushing the button on the OM1 shield using a Flush program (provided online), as required by the IACUC at our institution. Cages are changed every 2 weeks by replacing the DIY-NAMIC apparatus into a clean modified homecage. Modified cages with U-channels attached are compatible standard cage-washing procedures. Following each experiment in a box (when mice are removed), the plexiglass and enclosure are wiped down with babywipes and then with Clidox disinfectant spray. 70% ethanol is run through all water tubes and solenoids, and then allowed to dry. Every 4-6 months, or as needed, the noseports are removed from the plexiglass and enclosure for cleaning with soap and water. The tubing is replaced, and electronics are sprayed with compressed air to remove dust and dander, and wiped down with 70% ethanol when possible. A detailed cleaning protocol is available online.

### Drug administration

Following training as described above, and immediately following 3 days of exposure to the 9s delay in the test of impulsive action, mice were injected with drug or saline control. A selective 5-HT_1B_ agonist, CP 94253 hydrochloride (Tocris Bioscience Cat. No. 1317) was given at a high dose – 10 mg/kg, dissolved in 0.9% sterile saline and injected at 10ml/kg i.p (Knobelman et al., 2000). Injections were given within 30min of initiation of the dark cycle. Mice were randomly assigned to drug or vehicle conditions, and then 2-3 days after receiving their first injection, mice received the alternate condition. Data from the 12 hours (their dark phase) after receiving injections were analyzed.

### Statistical Analysis

Statistical testing was performed using analysis of variance (ANOVA), with post hoc Fisher’s least significant difference (LSD) in StatView (SAS Software, Cary, NC) or SPSS (IBM, Armonk, NY) for three-way ANOVAs. For the initial behavioral characterization, a repeated measures ANOVA was used to assess the change in behavior over days in the measures of number of rewards, number of ITI responses, and number of trials initiated per day. Post hoc Fisher’s LSD was used to assess differences between days. There were no significant effects of sex on any of the measures reported for the initial behavioral characterization (F_1,7_<3.0, p>0.05), though the sample size was not powered enough to rule out sex differences.

For the adolescent study, measures were summed over each day and then averaged across 3 days for each delay, except for proportion of correct trials which was averaged across each hour of the day, and then averaged across 3 days for each delay. A three-way repeated measures ANOVA was used to test the effect of sex and age on premature responding. There were no significant effects of sex on pokes during the delay window (F_1,14_=0.5, p=0.50), number of trial initiated (F_1,14_=3.3, p=0.0899), number of omissions (F_1, 14_=0.3, p=0.6159), or proportion of correct responses (F_1,14_=0.2, p=0.8947). There was an interaction of sex and delay length in the number of omissions (F_2,28_=5.3, p=0.0111) and proportions of correct responses (F_2,28_=3.8, p=0.0335), in that females had worse performance over all as the delay got longer, however there was no interaction with age and the pattern occurred in both adults and adolescents.. Two-way repeated measures ANOVAs, with post hoc Fisher’s LSD tests, were used to assess the effect of age on four behavioral measures over the 3 delay lengths [age (adol, adult) × delay length (3s, 6s, 9s)]: premature responding (nosepokes during the delay window), number of trials initiated, number of omitted trials, and proportion of correct trials.

For the pharmacology study, premature responses per trial was calculated by dividing the number of nosepokes during the delay period by the number of self-initiated trials for each hour, and a repeated measures ANOVA [condition (drug, saline) × time (1-12h)] was used to assess significance. All other measures were summed (number of delay responses, ITI responses) or averaged (proportion of correct responses and omissions), over 6h bins following injection (1-6h and 7-12h following injection) to analyze the effect of drug versus saline, based on drug half-life estimations from previous microdialysis studies following CP 94253 drug administration (Knobelman et al., 2000). In three-way ANOVAs (sex × drug × time) there were no significant main effects of sex on premature pokes during the delay (F_1,32_=0.1, p=0.8038), total trials initiated (F1,32=1.6, p=0.2129), ITI responding (F_1,32_=0.001, p=0.9913), or proportion of omissions (F_1,32_=0.2, p=0.6239). There were also no significant interactions of CP 94253 and sex on premature pokes during the delay (F_1,32_=0.8, p=0.3887), total trials initiated (F_1,32_=0.1, p=0.7704), ITI responding (F1,_32_=0.7, p=0.4099), or proportion of omissions (F_1,32_=0.2, p=0.7014). Although there was no main effect of sex on the proportion of correct trials (F_1,32_=2.6, p=0.1124), there was a significant interaction of sex and drug on proportion of correct trials, in that males that received drug had lower accuracy compared to females that received drug. Groups were then collapsed across sex, and repeated measures ANOVAs [condition (drug, saline) × time (1-6h, 7-12h)] were used to assess the effects of CP 94253 over the time, with post hoc Fisher’s LSD.

## Results

We designed, implemented, and tested the DIY-NAMIC system, and show that it is a low-effort, low-cost, and high-throughput method to measure complex behavior in mice (Fig 1). Slotted into a standard rodent home cage compatible with high-density racked cages, the DIY-NAMIC system allowed mice to rapidly learn complex operant behavioral tasks. Mice learn to nose poke at a cue-lit reward port to receive a liquid reward on a continuous reinforcement schedule (Fig 2A), receiving an average 172±13 rewards per day over the first three days of testing. Compared to the average 40-60 rewards/day in standard operant behavioral designs, this is a large increase in the number of trials/day that are run, therefore greatly speeding task acquisition. The majority of responses made were during the dark phase of their diurnal cycle; mice received only 10.6% of their rewards during the light phase (Fig 2B). This allows for more naturalistic testing of nocturnal rodents compared to many standard protocols which involve behavioral testing during the mouse dormant phase (daytime) for facilities that don’t have reverse light cycles. Additionally, intertrial interval (ITI) responding decreased by the second day (Fig 2C; F_2,16_=10.6; p<0.0012; D1vsD2: p=0.0008; D1 vs D3: p=0.0016; D2 vs D3: p=0.7359), showing that mice learned the stimulus-response association within a day.

Mice were subsequently trained to make an operant response (nosepoke) to an illuminated response-port (Fig 2D), and increased their earned rewards by the second day (Fig 2E; F_2,16_=4.6, p=0.0262; D1vsD2: p=0.0255; D1 vs D3: p=0.0136; D2 vs D3: p=0.7630), and further reduced ITI responding to the middle port by the third day of training (Fig 2F; F_2,16_=4.1, p=0.0372; D1vsD3: p=0.0378). Mice then learned to initiate trials (Fig 2G), and increased to stable performance by the second day (F_3,24_=5.7, p<0.0043; D1 vs D2: p=0.0054; D1 vs D3: p=0.0090; D1 vs D4: p=0.0007) self-initiating an average of 191±10 trials per day (Fig 2H). Following trial self-initiation training, mice learned to respond correctly on a discriminative cue paradigm, during which a response only in the illuminated port resulted in a reward (Fig 2I). A circadian rhythm of trials initiated was observed throughout 24h of testing with 89.9% of trials initiated in the dark phase (F_8,11_=12.6, p<0.0001). Within the dark phase, there was an effect of time on the number of trials initiated (F_8,5_=2.9, p=0.0269), with more trials initiated during the period 2 to 8 hours after lights off compared to other hours of the dark phase (Fig 2K). Mice performed correctly by responding with a nosepoke to the illuminated port on 84.7% of the 211.3±12.2 total trials, with incorrect responses on 13.8% of trials, and omissions on only 1.4% of trials (Fig 2J). The percentage of correct trials did not vary across the hours of the dark phase (F_8,5_=0.9, p=0.4734; Fig 2K).

Next, we assessed the capability of the DIY-NAMIC system to measure impulsive behavior using a modified 2-choice serial reaction time task (Fig 2L). Delays of varying lengths were introduced between trial initiation and the response cue in order to measure premature responding. The number of premature responses varied as a function of delay length (F_2,16_=5.6, p<0.0155; Fig 2M). Specifically, during the dark period (when mice initiated the majority of trials), there were more premature responses per trial during the 6 and 9 second delay, compared to the 3 second delay (3 vs 6: p=0.0260; 3 vs 9: p<0.0050; 6 vs 9: p=0.4357). Over days within each delay, the accuracy did not change (F_2,16_<1.53, p>0.05), however, omissions decreased across days during the 3 second delay (F_2,16_=14.3, p<0.0002; Fig 2N).

We next assessed impulsivity during adolescence, given that the adolescent period in mice is generally shorter than the time needed for training on many traditional operant paradigms. While our goal was to measure impulsivity, it was important to also obtain additional measures of attention, motivation, and performance. Compared to adults, adolescent mice showed increased impulsive behavior as measured by premature responding (Fig 3A; F_1,16_=8.54, p=0.0099) with more premature responding in adolescents as the delay got longer (F_2, 32_=24.9, p<0.0001). Specifically, adolescents had significantly higher levels of responding during the 9 second delay period (p=0.0491 for 3s; p=0.0521 for 6s; p=0.0281 for 9 sec). This suggests that premature responding is a readout of the adolescents’ reduced ability to inhibit responding since the number of responses increases as the difficulty/delay increases, rather than a reflection of their observed general hyperactivity or increased motivation. Importantly, adolescents did initiate more trials (Fig 3B; F_1,16_=5.8, p=0.0280), however this did not vary with the change in delay/difficulty of the paradigm (F_2,32_=0.2, p=0.8162). We also used the number of omitted trials as a readout of attention. Adolescent mice showed decreased attention as measured by increased number of omission trials (Fig 3C, F_1,16_=7.1, p=0.0173). Again, there was no effect of delay length on omissions suggesting that the decreased attention was unlikely the cause of the increased impulsivity (F_2,32_=1.2, p=0.7256). Importantly, adolescents did not show differences in the proportion of correct trials (Fig 3D, F_1,16_=0.1, p=0.7680) indicating that their performance on the task was normal and there was no effect of age on accuracy or overall performance on the paradigm. Overall, the results suggest that adolescent mice display increased impulsive action as measured by premature responding. Furthermore, these data demonstrate the ability to test complex self-initiated operant behaviors during limited time frames, such as adolescence, in the DIY-NAMIC system.

**Figure 3.**
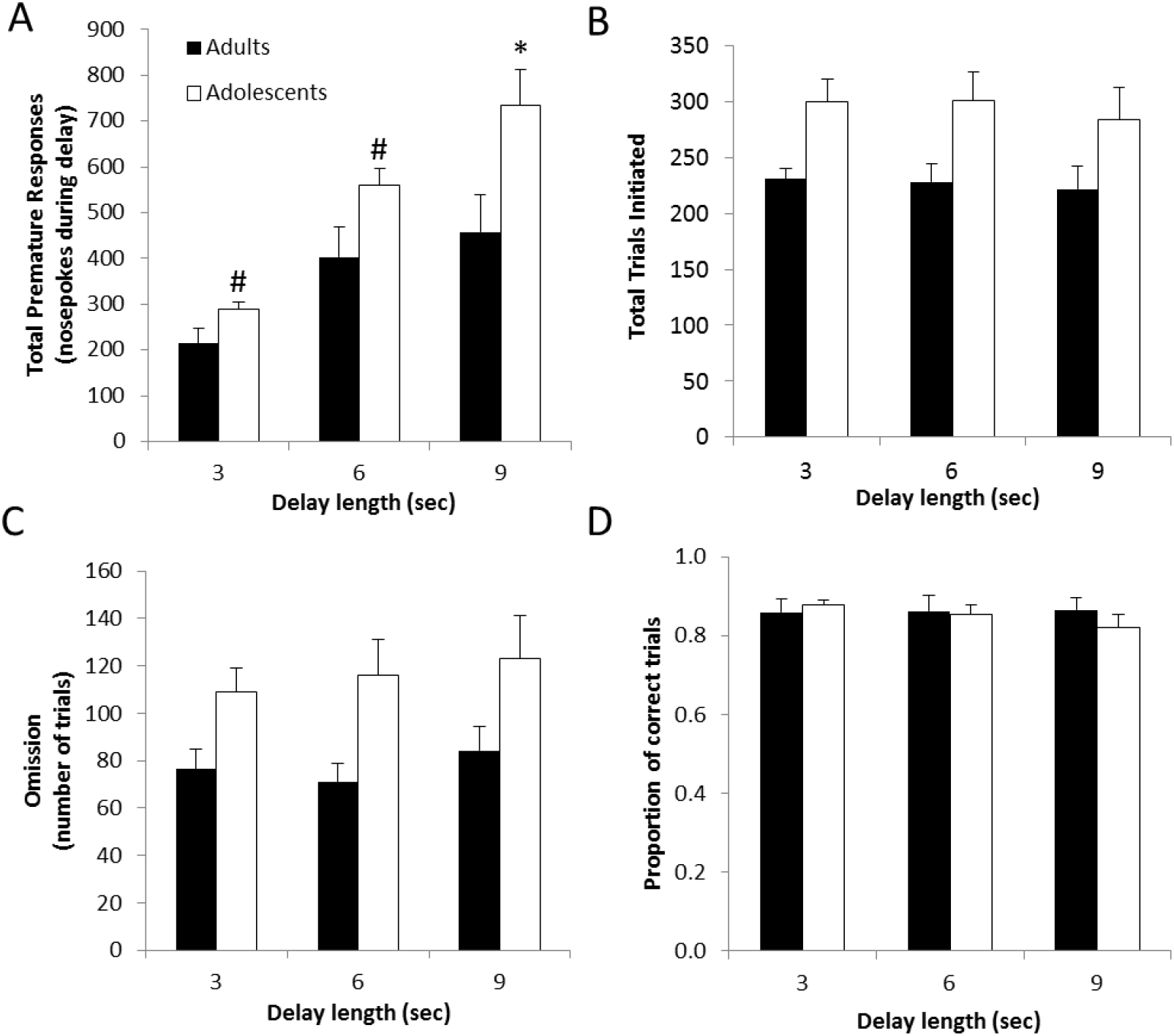
Adolescents show higher levels of impulsive behavior. Premature responses as measured by total nosepokes during the delay period (A), total initiated trials (B), average number of omitted trials (C), and proportion of correcttrials of those attempted (D) are shown for adults and adolescents for the three delay lengths. Group averages are shown for each delay averaged over three days, with error bars representing the SEM; *, p<0.05, #, p=0.05.

Finally, in a pharmacology experiment, we tested the effect of manipulating serotonin signaling on impulsive action using DIY-NAMIC boxes. We focused on the effect of activating the 5-HT_1B_ receptor by administering the receptor-specific agonist, CP 94253, and measuring behavior using a modified 2-choice serial reaction time task in the DIY-NAMIC cages. Compared to operant behavior which traditionally measures behavior at single point in time, the DIY-NAMIC system allowed investigations into the timecourse of drug effects on behavior (Fig 4A, main effect of time: F_11,374_=3.3, p=0.0002). Administration of the agonist resulted in decreased impulsivity, measured by reduced premature responding, which varied over time as the drug effects dissipated (Fig 4B; interaction of drug × time: F_1,34_=4.8, p=0.0352). Specifically there was decreased responding during the delay period in the 6h following injection of the 5-HT_1B_ receptor agonist (p=0.0147), and not during the subsequent 6h of the dark phase (p=0.4595). Importantly, the agonist also decreased the total number of trials initiated in the first 6h after drug administration, suggesting a decrease in general motivation (Fig 4C, interaction: F_1,34_=10.6, p=0.0026; drug vs vehicle for 1-6h: p=0.0141). While this is a relevant factor in interpreting the premature responding, the correction for premature responses per trial (Fig 4A), suggests that the drug effects on impulsive responding is not solely driven by decreases in motivation. Additionally responding during the ITI can be driven by both impulsive and general hyperactive behavior; these responses were slightly, but not significantly decreased for the first half of the dark cycle following drug treatment (Fig 4D, interaction drug × time: F_1,34_=3.1, p=0.0873). Finally, there was also a significant effect of the agonist on the proportion of omitted trials (Fig 4E; F_1,34_=6.3, p=0.0170), and this was only significant for the first 6h following drug administration (interaction of time × drug: F_1,34_=4.5, p=0.0412). There was no effect of the agonist on the proportion of correct trials of those attempted (Fig 4F; F_1,34_=2.0, p=0.1670), indicating the drug effects were specific to premature responding, and likely not due to effects on performance or attention. These data show the utility of DIY-NAMIC boxes in assessing pharmacological manipulations, particularly in allowing for time-course data. Additionally, these data show a novel effect of serotonin pharmacology on impulsivity.

**Fig 4.**
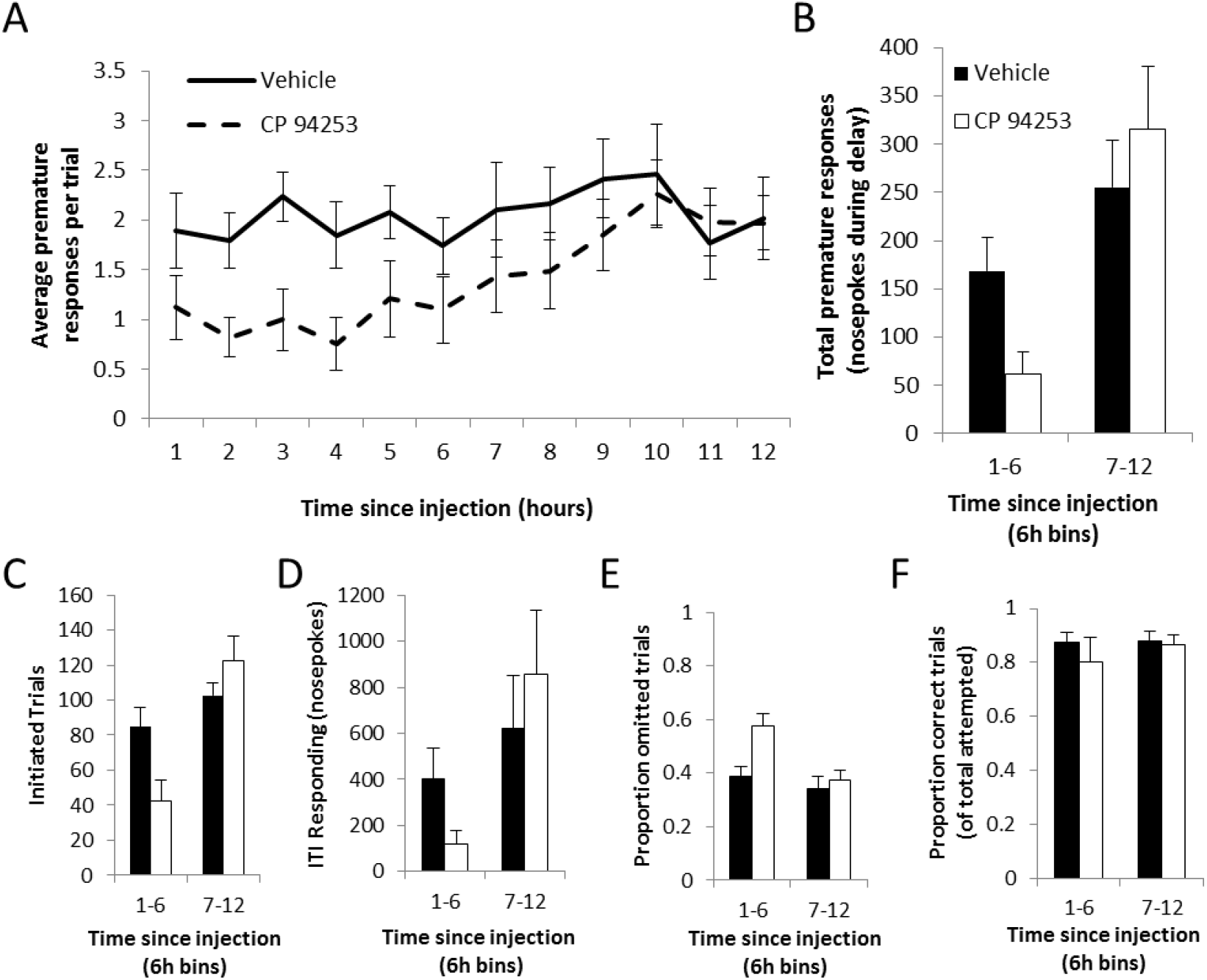
5-HT_1B_ agonist CP 94253 reduces impulsivity. (A) DIY-NAMIC boxes allow for continuous assessment of the behavioral effects of pharmacological manipulations on impulsive action. The average number of nosepokes during the 9s delay window per trial is shown over the 12h dark phase immediately following injection of vehicle or drug (CP 94253). (B) Binned by 6h, premature responses are shown for the first and second halves of the dark phase following vehicle or drug administration. Other measures includingtotal number of trials initiated (C), ITI responding (D), proportion of omitted trials (E), and proportion of correct attempted trials (F) are shown for drug and saline conditions over first and second half of dark phase.

## Discussion

Given the recent push to explore more robust and multi-dimensional behavioral readouts of animal models for neuropsychiatric disorders, the development of high-throughput and automated behavioral screens are critical (Gordon and Dzirasa, 2016; Bale et al., 2019). Operant behavioral paradigms provide rich behavioral datasets, however they are labor intensive and lengthy (daily training for weeks/months), and generally require food deprivation and costly equipment. To address these limitations, we developed an Arduino-based system as a homecage, low-touch, fully automated operant chamber that allows mice to self-initiate trials for their daily water intake. Data is logged continuously to a PC, and experimenter effort is only required to monitor/weigh animals to assess welfare, and upload new programs to Arduinos when switching behavioral paradigms (though even this could also be automated in future implementations). We have shown that mice learn basic Pavlovian and operant behaviors over a few days in DIY-NAMIC cages, performing hundreds of trials in a day in their dark/active phase. Additionally we have validated its use in measuring effects of drugs on behavior, allowing for monitoring over long time-frames. Finally, we show that we can measure impulsive behavior in adolescent mice, and show increased impulsivity compared to adults.

The difficulty of training mice on complex operant behavioral paradigms within a few weeks using standard procedures and equipment limits their utility in examining adolescent behavior in mice given its short timeframe (Workman et al., 2013; Brust et al., 2015). We have found this to be a critical limiting factor in understanding the adolescent development of the neural and behavioral basis of impulsivity, an important sensitive period for the etiology and pathogenesis of neuropsychiatric disorders which include impulsive behavior as a core phenotype (Chambers et al., 2003; Verdejo-Garcia et al., 2008). Our studies reported here measure behavior in mid-to late-adolescence, though the DIY-NAMIC system easily allows future studies to characterize impulsivity throughout various stages of adolescence including early adolescence (~30-40 days).

There have been past efforts to study operant behavioral measures in adolescent mice. Some of these studies have taken the strategy of eliminating mice that don’t learn quickly enough to progress to increasingly complex behavioral paradigms, leading to potentially biased samples (Arnold and Newland, 2018). There has also been some progress in the development of new training paradigms that reduce training time to allow testing in adolescent mice (Ciampoli et al., 2017; Sasamori et al., 2018), however they still require food deprivation which could be detrimental to growth and the maturation of feeding circuits during adolescence. Inexpensive Arduino-based operant testing apparatus have also been developed, such as ROBucket (Devarakonda et al., 2016), though these have also been used with the daily experimenter-initiated daily sessions. Finally, others have used modified commercially available designs such as the CombiCage which allow free-access to the standard costly operant conditioning chambers via a tube connected to a homecage to measure adolescent impulsive action (Remmelink et al., 2017). Overall, the DIY-NAMIC system provides an improved method to measure impulsivity in adolescent mice by combining the benefits of some preceding technology and allowing testing in a number of behavioral domains with minimal experimenter effort and high-throughput protocols.

We were able to use the rich data set collected from the DIY-NAMIC system to tease impulsivity apart from other behavioral parameters that may also be different during adolescence and could otherwise obscure or contaminate measures of impulsive behavior. For example, previous work has shown hyperactivity, increased motivation, and decreased attention in adolescent mice (Ciampoli et al., 2017). We were able to increase the sensitivity of the task to detect impulsivity by increasing the delay length (Dalley et al., 2002). Our measure of impulsivity increased as delay length increased, while measures of hyperactivity, motivation, and attention did not. Interestingly, our results differed from a previous study which used a CombiCage for 24h access from the homecage to a commercial testing chamber and reported no significant differences between adult and adolescent mice in a 5-CSRTT (Remmelink et al., 2017). Although there were a number of methodological differences between our studies, including the use of food pellets for rewards and a paradigm which seemed to result in many more initiated trials with a higher percentage of omitted trials. This may suggest a more challenging task which may have obscured age differences. On the other hand, our results were consistent with another previous report of impulsivity measured in adolescent mice using a modified training procedure for 3-CSRTT in a traditional session-based design in commercial equipment (Sasamori et al., 2018). These authors report increased impulsive action in late adolescent mice as measured increased by premature responses, however did not see increased omissions as seen in our data.

Automated home cage phenotyping provides the opportunity to measure behavior in a more reproducible manner because it limits experimenter-introduced variability (Bains et al., 2018). A system that is compatible with high-density racked cages and does not include video-based phenotyping (because of the long time-frames) provides the most usable and reliable method. The DIY-NAMIC system also provides a data-rich, low-effort approach to psychoactive drug screening. We show here that the effects of acute administration of drugs can be assessed over prolonged periods of time. This is an important improvement compared to many “single-use” behavioral tests which can only reliably be performed once, requiring higher numbers of animals in order to be able to look at the temporal profile of drugs. Additionally, the DIY-NAMIC system provides an opportunity to assess longer-term effects of acute administration, as well as effects of chronic drug administration, with lower effort and animal numbers than traditional testing. Finally, the shortened training time required allows for the preclinical testing of drugs in adolescence, which is important given potential differential actions of drugs over the lifecourse.

Homecage behavioral testing allows for behavioral analysis across the circadian cycle, an important variable for the interpretation of studies assessing impulsivity and other motivated behavior, that is often overlooked (Balachandran et al., 2020). Conventional session-based, non-homecage behavioral testing generally occurs during the light phase of the circadian rhythm, likely causing partial entrainment to feeding occurring in the light cycle, and resulting in dysregulation of homeostatic systems (Gritton et al., 2012). Furthermore, it is difficult to test the effect of circadian rhythms using traditional session-based operant paradigms which is critical because many manipulations, including drug administration, have interactions with time-of-day. On the other hand, the lack of control over the timing of trials can also create a confound in the interpretation of data collected over the circadian cycle. Since mice are more active/motivated for water in the dark phase, assessing the effect of circadian period on behaviors is difficult. Behavior can be normalized by trials initiated, though in many hours during the light phase there are very few, or no, trials initiated. An alternative option with this system would be, following acquisition of task behavior, to restrict access to short (1h) timeframes by only allowing trial initiation during “session-like” time periods. This may also be a necessary approach if using manipulation, recording, or imaging techniques to force trials within a limited time period.

Although the DIY-NAMIC system provides a number of significant advantages over standard commercially available operant boxes, it is not without its own disadvantages. One downside currently is the need to single-house mice. Although the daily stressor of handling is eliminated, singly-housed mice may also result in effects on baseline stress levels and/or behavioral readouts (Meijer et al., 2007; Kamakura et al., 2016; Hebda-Bauer et al., 2019). This limits the interpretation of some results, for example, the differences we see in adolescent impulsive behavior could be in part due to a differential response to single-housing. The DIY-NAMIC system is compatible with standard homecage enrichment, and we include nesting material and an igloo hut in all cages. Future iterations of this system could use an RFID chip implanted subcutaneously, and a reader in an access port to the receptacles to allow group housing. Another technical improvement is the use of WiFi/Bluetooth microprocessor board and shield (see OpenMaze.org) to allow wireless data upload/program download. Although our facility allows USB/power wires from the cages on high-density racks, some facilities or IACUCs may not allow this and the use of the wireless data transmission and onboard batteries could eliminate wiring connections completely.

We used the DIY-NAMIC system here to measure impulsive action, but many other paradigms could be implemented using this method including random ratio paradigms and other types of impulsivity, such as delayed discounting. However, paradigms that require manipulation of context would not be possible in this system. Given the Arduino platform and the modular build of the DIY-NAMIC system, additional stimuli and cues (e.g. vibration pad for tactile stimulus) would be necessary customization changes in order to measure some addtional phenotypes.

In conclusion, we have developed the DIY-NAMIC system as a low-touch, low-cost homecage operant behavioral testing system with an opensource platform. All materials are commercially available, and build instructions are documented online in detail, including tips for researchers with minimal electrical and material engineering skills. Software for all of the programs and analysis used for experiments in this manuscript are also available online, including Arduino programs for running behavioral paradigms, Processing scripts for logging data, and Python scripts for data analysis and plotting. The DIY-NAMIC system greatly increases the ability to measure complex and robust behavioral measures in a high-throughput manner, enabling more productive and reproducible basic research in animal models for neuropsychiatric disorders.

